# Translation Inhibition by Rocaglamide A Enhances Susceptibility of Yeasts to Caspofungin

**DOI:** 10.1101/2025.09.21.677667

**Authors:** Qian He, Peter J. Schuessler, Aravind Srinivasan, Jessie Caprino, Joseph Barbi, Sarah E. Walker

**Affiliations:** Department of Biological Sciences, SUNY at Buffalo, Buffalo, New York, 14260, USA; Department of Immunology, Roswell Park Comprehensive Cancer Center, Buffalo, New York, 14263, USA; Department of Thoracic Surgery, Roswell Park Comprehensive Cancer Center, Buffalo, New York, 14263, USA

**Author notes:** Address correspondence to Sarah E. Walker.

## Abstract

Fks1 catalyzes the synthesis of β-1,3-glucans, a major structural component of the fungal cell wall, and is the primary target of the antifungal drug caspofungin. Mutations in Fks1 confer caspofungin resistance by disrupting its interaction with the drug, thereby reducing inhibition of Fks1 enzymatic activity. Previous studies demonstrated that translation of the *FKS1* mRNA was highly dependent on the translation initiation helicases eIF4A and Ded1 (1). Therefore, we investigated whether treatment of *Saccharomyces cerevisiae* cells with the eIF4A inhibitor Rocaglamide A (RocA) or mutation of Ded1 affects translation of *FKS1* and susceptibility to caspofungin. Using WT and temperature-sensitive *ded1-ts* strains, we demonstrated that RocA enhanced caspofungin-mediated growth inhibition and translation repression. Sensitivity to both drugs was further enhanced in *ded1-ts* strains, suggesting specifically targeting Ded1 in fungi could be an effective mechanism to prevent caspofungin resistance. We extended the analysis to *Candida glabrata*, a related fungal pathogen, and found similar results. Importantly, combining RocA with caspofungin was fungicidal in both species, suggesting the combination could decrease development of caspofungin resistance in pathogenic yeasts. Together these findings highlight the potential of targeting translation initiation helicases for effective combination antifungal treatments.

## INTRODUCTION

The fungal cell wall is essential for viability, and defects in its synthesis typically result in loss of structural integrity and cell death. Importantly, the fungal cell wall is composed of components such as chitin, glucan, and mannan, which are absent in humans, making it an attractive target for antifungal drug development (2).

Among the major cell wall components, β-1,3-glucan is universally present in fungal species and plays key roles in maintaining cell shape, forming a physical barrier, and serving as a scaffold for the attachment of other wall elements (3). Its synthesis in nearly all fungi is catalyzed by a β-1,3-glucan synthase Fks1, encoded by the *FKS1* gene in *S. cerevisiae* (4). This protein is not present in mammalian cells, making it an attractive target for antifungals. Importantly, Fks1 is the target of caspofungin, the first clinically-approved echinocandin. This drug displays broad-spectrum inhibitory activity against *Candida* species (5), but the emergence of resistant strains has raised concerns about long-term utility. Since mutations in *FKS1* confer caspofungin resistance (5, 6), strategies to target Fks1 expression could inform efforts to achieve optimal clinical use of the drug.

Translation inhibition is a major mode of action for bacterial antibiotics, with several classes of drugs that target the ribosome and elongation factors. Similar translation elongation drugs have also been found effective in combatting fungal growth, either alone or in combination with other drugs (7-9). More recently, a class of eukaryotic translation initiation factor inhibitors, the rocaglates, was demonstrated to be effective in preventing growth of various budding yeasts, and in particular the emerging pathogen *Candida auris* (10). Rocaglates clamp translation initiation helicases eIF4A and DDX3 to mRNA, blocking ribosomal preinitiation complex scanning to the start codon (11).

In this study, we investigated the ability to impair expression of Fks1 and other proteins at the level of translation initiation as a means to heighten caspofungin susceptibility. We found that when combined with caspofungin, RocA enhanced its growth inhibition and reduced translation of an *FKS1* translation reporter. Moreover, a *ded1-ts* mutant was even more susceptible to drug treatment, suggesting targeting Ded1 specifically could be an effective antifungal tactic. To test whether these observations extend to clinically-relevant fungi, we examined *C. glabrata*, a close relative of *S. cerevisiae* ranked fifth on the 2022 World Health Organization fungal priority pathogens list (12). *C. glabrata* showed higher intrinsic resistance to caspofungin than *S. cerevisiae*, but similar sensitivity to RocA. We found that the combination of RocA and caspofungin restricted growth of *C. glabrata* to a greater degree than either drug alone, and this effect was fungicidal. Altogether, these results suggest that preventing the expression of a drug target via translation inhibition enhanced drug activity, which could be a promising strategy for combination antifungal therapies.

## MATERIALS AND METHODS

### Construction of yeast strains and plasmids

All strains, plasmids, and primers used in this study are listed in Tables 1, 2, and 3, respectively. Plasmid pJC2 was constructed by QuikChange using primers JC9 and JC10 and pCAS as the template. pJC3 and pJC5 were generated by Gibson assembly using BamHI-digested pRS416 and pRS415 and PCR products amplified with Phusion polymerase and primers JC18, JC19, JC23, and JC24, with BY4741 genomic DNA as the template (NEB #E2611L, R0136L, and M0531L). Plasmid pPS9 was constructed using the QuikChange Lightning Multi Site-Directed Mutagenesis Kit (Agilent #210514) with primers PS45-PS48 and pJC5 as the template. Plasmids pPS7, pPS8, pPS10, and pPS11 were constructed by Gibson assembly: pPS7 used PCR fragments from primers PS39 and PS40 with pEKD1024 as template and primers PS41 and PS43 with BY4741 genomic DNA as template; pPS8 used a fragment amplified from pPS7 with primers PS49 and PS50 and BamHI- and EcoRI-digested pHO-hisG-URA3-hisG-poly-HO (Addgene #51663); pPS10 plasmid used fragments amplified from pPS8 using primers ND5 and ND6, and from BY4741 genomic DNA using primers ND7 and ND8; and pPS11 used a fragment amplified by primers PS51 and PS52 from pPS10 with BglII-digested pHO-Poly-KanMX4-HO (Addgene #51662).

**Table 1:** Yeast strains used in this study Strain Name Genotype.

| Strain Name | Genotype |
| --- | --- |
| YSW3 | *MATa, his3Δ1, leu2Δ0, ura3Δ0, met15Δ0, TIF3* |
| YSW146 | *MATa, his3Δ1, leu2Δ0, ura3Δ0, met15Δ0, ded1Δ0 [pJC3]* |
| YSW193 | *MATa, his3Δ1, leu2Δ0, ura3Δ0, met15Δ0, ded1Δ0, [pPS09]* |
| YSW194 | *MATa, his3Δ1, leu2Δ0, ura3Δ0, met15Δ0, ded1Δ0, [pJC5]* |
| YSW222 | *MATa, his3Δ1, leu2Δ0, ura3Δ0, met15Δ0, hoΔ0::PGAL10-mCherry-PGAL1-FKS1 5' UTR-GFP ded1Δ0 [pPS09]* |
| YSW223 | *MATa, his3Δ1, leu2Δ0, ura3Δ0, met15Δ0, hoΔ0::PGAL10-mCherry-PGAL1-FKS1 5' UTR-GFP ded1Δ0 [pJC5]* |
| FSW114 | *Candida glabrata BG2 (Wild-type)* |

**Table 2:** Plasmids used in this study.

| Addgene ID | Plasmid Name | Relevant Gene | Markers | Parental Vector | PCR Template | Primers | Source |
| --- | --- | --- | --- | --- | --- | --- | --- |
| 60847 | pCas | Cas9 with sgRNA | *E. coli* KanR, Yeast KanMx |  |  |  | (19) |
| N/A | pRS415 | Yeast centromere vector | *E. coli* AmpR, Yeast *LEU2* |  |  |  | (20) |
| N/A | pRS416 | Yeast centromere vector | *E. coli* AmpR, Yeast *URA3* |  |  |  | (20) |
| N/A | pEDK1024 | GFP:mCherry bidirectional reporter without insert upstream of GFP | *E. coli* AmpR, Yeast *MET15* |  |  |  | (13) |
| 51663 | pHO-hisG-URA3-hisG-poly-HO | Empty vector for integration of target sequence at HO locus | *E. coli* AmpR, Yeast *URA3* |  |  |  | (21) |
| 51662 | pHO-Poly-KanMX4-HO | Empty vector for integration of target sequence at HO locus | *E. coli* AmpR, Yeast *KanMx* |  |  |  | (21) |
| TBD | pJC2 | sgRNA targeting *DED1* | *E. coli* KanR, Yeast *KanMx* | pCas | N/A | JC9/10 | This study |
| TBD | pJC3 | WT Ded1 Silent mutation | *E. coli* AmpR, Yeast *URA3* | pRS416 | BY4741 DNA | JC18/19/23/24 | This study |
| TBD | pJC5 | WT Ded1 Silent mutation | *E. coli* AmpR, Yeast *LEU2* | pRS415 | pJC3 | JC18/19/23/24 | This study |
| TBD | pPS09 | Temperature sensitive mutant ded1 (W253R, T408I) | *E. coli* AmpR, Yeast *LEU2* | pJC5 |  | PS45/46/47/48 | This study |
| TBD | pPS07 | GFP:mCherry bidirectional reporter with *SLK19* 5' UTR | *E. coli* AmpR, Yeast *MET15* | pEDK1024 | BY4741 DNA | PS39/40/41/43 | This study |
| TBD | pPS08 | GFP:mCherry<br>bidirectional<br>reporter with<br>*SLK19* 5'<br>UTR | *E. coli* AmpR,<br>Yeast *URA3* | pHO-hisG-URA3-hisG-poly-HO | pPS07 | PS49/50 | This<br>study |
| TBD | pPS10 | GFP:mCherry<br>bidirectional<br>reporter with<br>*FKS1* 5' UTR | *E. coli* AmpR,<br>Yeast *URA3* | pPS08 | BY4741<br>DNA | ND5/6/7/<br>8 | This<br>study |
| TBD | pPS11 | GFP:mCherry<br>bidirectional<br>reporter with<br>*FKS1* 5' UTR | *E. coli* AmpR,<br>Yeast *KanMx* | pHO-Poly-KanMX4-HO | pPS10 | PS51/52 | This<br>study |

**Table 3:**
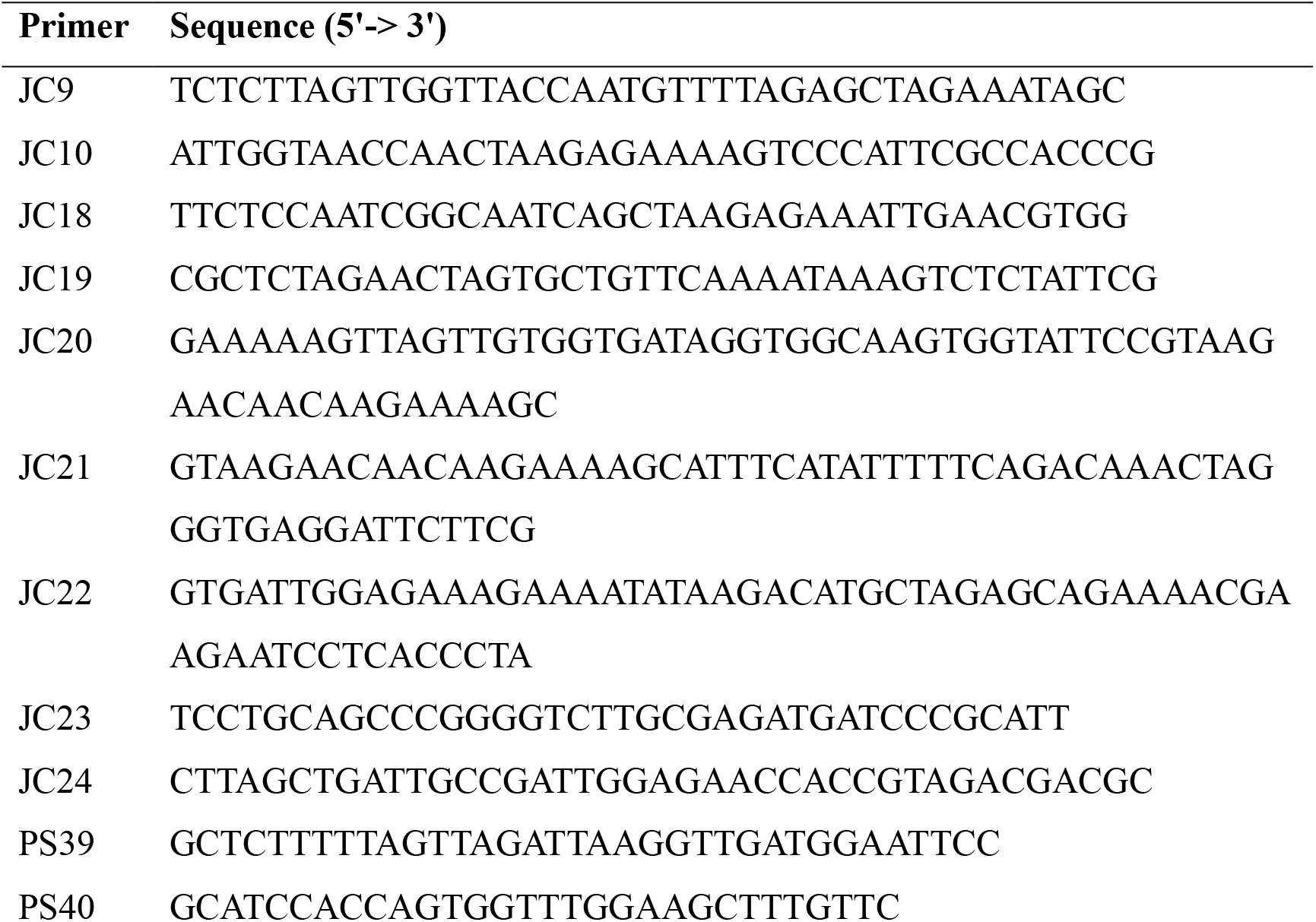
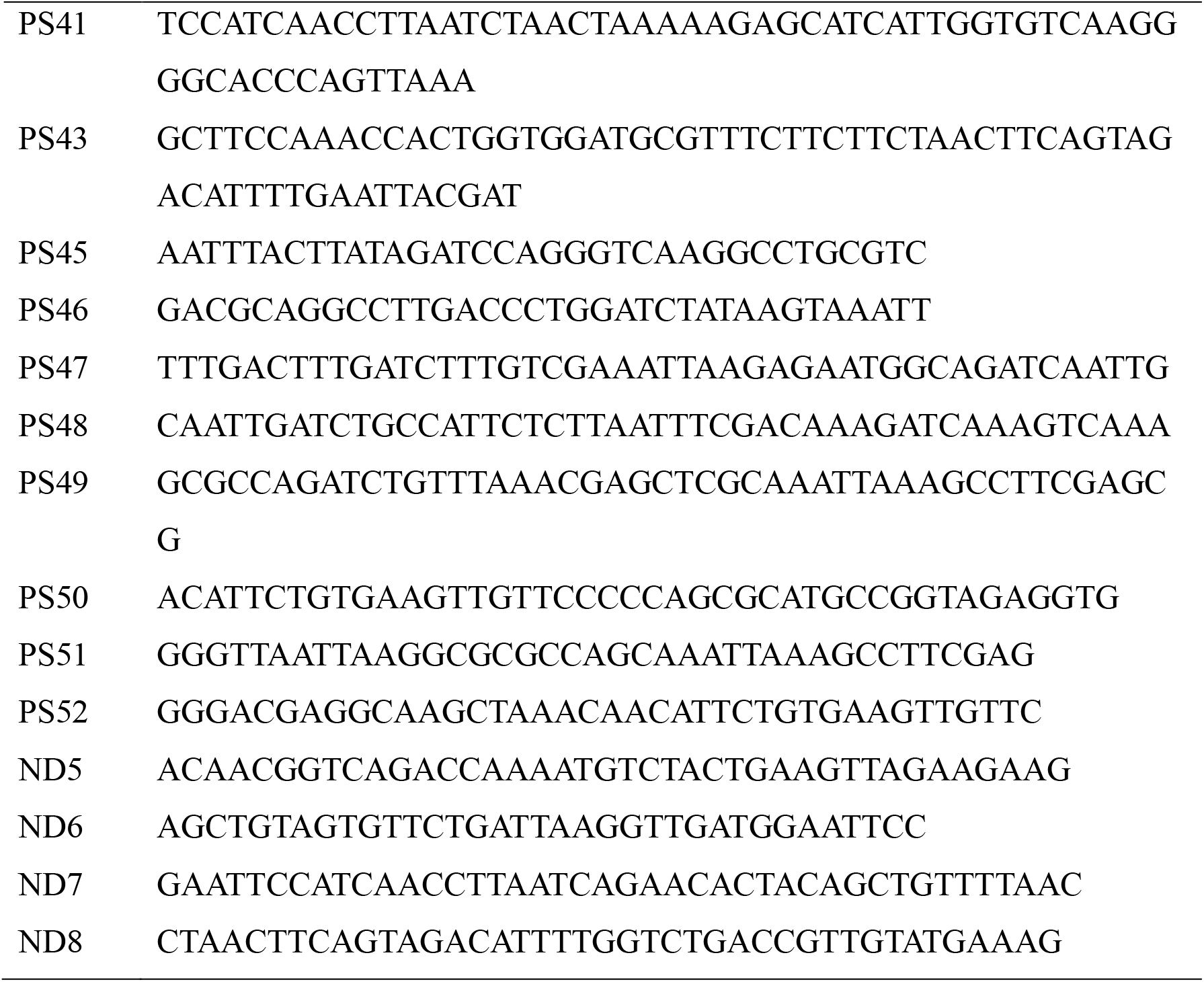
Primers used in this study.

A strain of *S. cerevisiae* lacking a chromosomal copy of *DED1* (YSW146) was generated by CRISPR/Cas9. Here, the parent strain (BY4741) was transformed with the plasmid pJC2, encoding Cas9 and an sgRNA targeting *DED1*, pJC3 encoding *DED1* with silent mutations that eliminated the corresponding sgRNA binding site alongside a counter-selectable URA3 marker, and a homology-directed repair template to eliminate the *DED1* open reading frame, followed by selection on G418-containing media. The homology-directed-repair template fused the 5′ and 3′ untranslated regions (UTR) of *DED1* and was generated by Phusion PCR with 3 overlapping oligos (JC20, JC21, and JC22). Following colony-PCR screening and sequencing, isolates were grown without G418 and loss of G418-resistance was confirmed. WT and *ded1-ts* strains (YSW194 and YSW193) were then generated by plasmid shuffling with LEU2 plasmids pJC5 and pPS09. YSW194 and YSW193 were then transformed with the Not1-digested 3 kb fragment of pPS11 to integrate the *FKS1* reporter cassette, and transformants were selected on YPD media with G418 to obtain YSW223 and YSW222.

### Chemical susceptibility and recovery assays

Single isolates of strains freshly-streaked from freezer stocks were grown overnight in SC-raffinose media (2% raffinose plus 0.1% dextrose as carbon sources) at 30°C with shaking and then diluted to an OD_600_ of 0.035 in SC-galactose media (2% galactose plus 2% raffinose) to induce expression of the dual-fluorescence reporter. *Candida glabrata* BG2 (gift from Laura Rusche) was cultured overnight in YPD media and similarly diluted to an OD_600_ of 0.01 in fresh YPD. Compound potency was evaluated by adding indicated concentrations of compounds to 96-well plates. OD_600_, GFP, and mCherry signals were recorded every 30 minutes for 48 hours at 37°C using a Spark plate reader (Tecan, Switzerland) with double-orbital shaking. For *C. glabrata*, cells were grown at 37°C and only OD_600_ was measured. All data were corrected for background signals from media-only controls. At least three biological replicates were performed, typically also including two technical replicates.

After 48-hours growth in the chemical susceptibility assay, cells were spotted onto YPD agar plates lacking drugs to assess the fungistatic or fungicidal nature of the indicated agents. Plates were incubated at indicated temperatures and photographed after 24 hours.

### Flow cytometry analysis

The sensitivity of the *FKS1* fluorescent reporter to Ded1 inhibition was confirmed by flow cytometry. Here, strains carrying wild type *DED1* (YSW223) or the temperature sensitive *ded1-ts* mutant (YSW222) on the *FKS1* reporter plasmid were incubated overnight at either 30°C or 37°C, with and without induction of the reporter. After washing and resuspending cells in sterile phosphate buffered saline, proportions of single cells expressing GFP, mCherry, or both signals were assessed with a BD LSR II multi-color flow cytometer housed in the Flow and Immune Analysis Shared Resource at Roswell Park Comprehensive Cancer Center. Non-reporter strains and uninduced cells served as negative controls, while strains expressing a single fluorescent protein were used to define populations during analysis carried out using Flowjo Software.

## RESULTS

### Preliminary assessment of compound sensitivity in WT and *ded1-ts* strains

Previous work analyzed transcriptome-wide differences in translation by ribosome profiling when translation initiation factor helicases eIF4A and Ded1 were mutated in yeast (1). Among the RNAs showing significant reductions in translation when either factor was depleted was the *FKS1* transcript, encoding the Fks1 β-1,3-glucan synthase targeted by caspofungin. This prompted us to ask whether inhibition of eIF4A and/or Ded1 would reduce expression of Fks1 protein, thereby minimizing the amount of caspofungin needed to inhibit growth.

To test this, we employed the translation initiation inhibitor Rocaglamide A (RocA), a member of the rocaglate family that impedes both eIF4A and the Ded1 ortholog, DDX3, in cancer by promoting clamping of the helicases to polypurine motifs, which then blocks ribosomal scanning (11). We hypothesized that treatment with RocA would affect both eIF4A and Ded1, reducing Fks1 expression to augment caspofungin. We also tested effects of both drugs on a strain (*ded1-ts*) harboring temperature-sensitive mutations of Ded1 shown to confer large reductions in Fks1 translation (1).

We first conducted chemical susceptibility assays for each compound in isolation. WT and *ded1-ts* growth was monitored in liquid media for 48 hours at 30°C. Next, cells were plated on media lacking drug to assess whether the cells were able to recover from compound exposure at three temperatures: 30°C is permissive for *ded1-ts* while growth of the mutant is inhibited at 36°C and 37°C due to partial inactivation of Ded1.

We found that 0.08 μg/ml caspofungin substantially inhibited growth of WT and *ded1-ts* strains, consistent with a reported minimum inhibitory concentration (MIC) of caspofungin for *S. cerevisiae* of 0.09 μg/ml (6). A dose-dependent growth defect was observed for caspofungin in both strains (Fig. 1A). Similarly, RocA caused a dose-dependent reduction in growth in both WT and *ded1-ts* strains at concentrations between 120 and 500 μM (Fig. 1B). The similar performance of WT and mutant strains in this assay is consistent with the nature of the *ded1-ts* mutation, which does not affect growth at 30°C.

**Figure 1.**
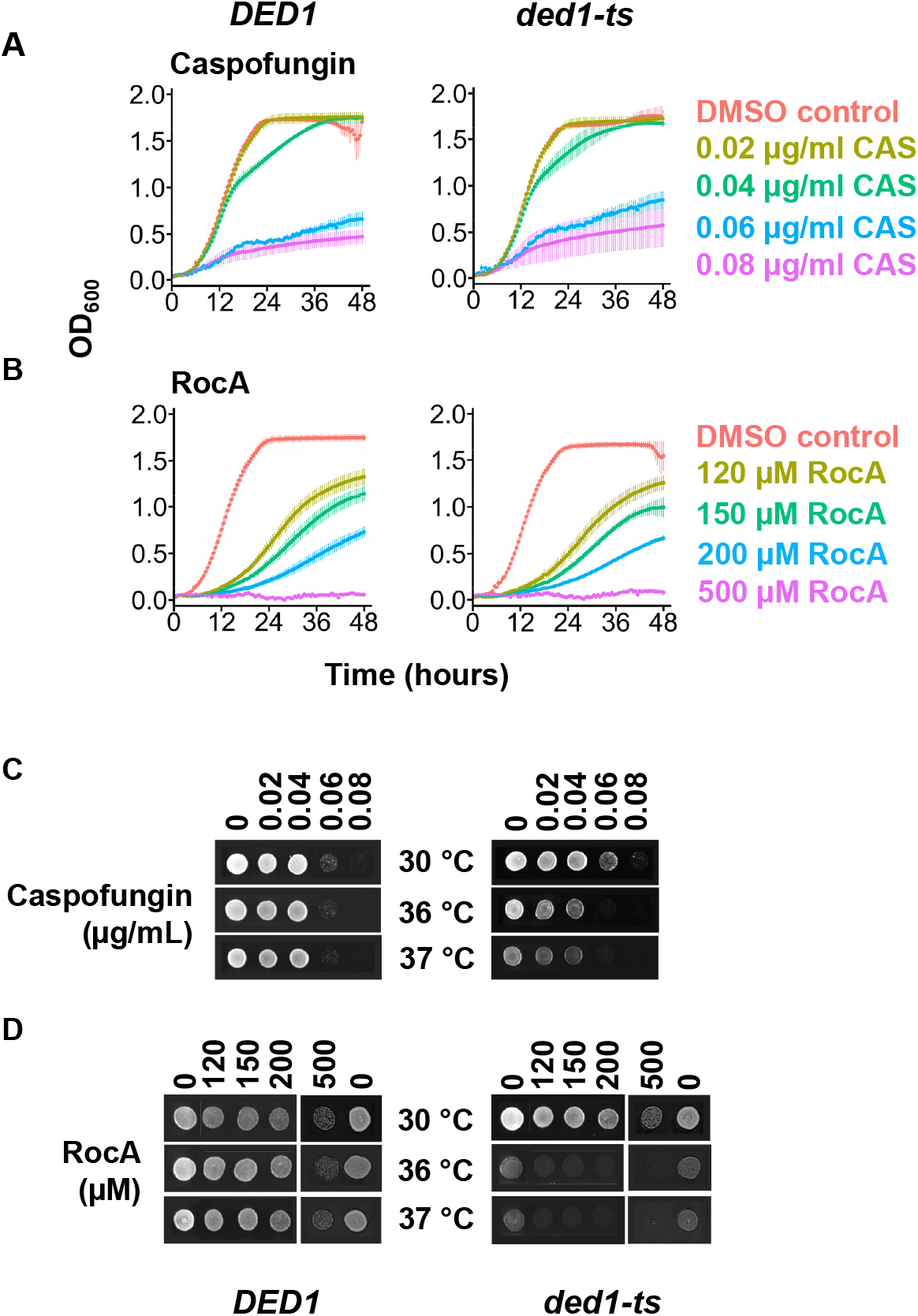
Assessment of *S*. cerevisiae susceptibility to caspofungin and RocA and recovery from treatment as a function of Ded1 activity. **A-B**. 48-hour growth curves of *DED1* (YSW223) and *ded1-ts* (YSW222) strains treated with increasing concentrations of caspofungin (CAS, A), or RocA (B) at 30°C. Data represents the mean± standard deviation of the mean from 3 biological replicates with 2 technical replicates. **C-D**. Recovery following compound exposure: *DED1* and *ded1-ts* cells were spotted on YPD plates and incubated for one day at the indicated temperatures following 48 hours treatment with indicated compounds.

In recovery assays, prior treatment with 0.08 μg/ml caspofungin prevented recovery of both WT and *ded1-ts* strains. As expected, growth of the *ded1-ts* strain was generally impaired at higher temperatures (Fig. 1C). In addition, *ded1-ts* strains exhibited further defects when treated with RocA and recovered at higher temperatures, compared to untreated controls, consistent with RocA and the *ded1-ts* mutation targeting the same process (Fig. 1D). This analysis validated the optimal concentrations of RocA and caspofungin for analyzing combinations of the drugs, e.g. 0.06 μg/ml or lower caspofungin, and 500 μM or lower RocA.

### RocA and caspofungin showed synthetic effects on growth and translation

In order to monitor translation effects of the drug combinations, we used a dual-fluorescent reporter derived from the previously described RNA-ID reporter, in which the 5′ UTR and first 30 nucleotides of the *FKS1* gene were fused to Superfolder-GFP under control of a bidirectional galactose promoter (13). When cells are grown in galactose-containing media, this system also induces transcription of an mCherry reporter RNA with a short 5′ UTR, predicted to be less structured and less dependent on Ded1 activity (Fig. 2A). We first determined the effects of Ded1 activity on the expression of the FKS1-GFP and mCherry control reporters by flow cytometry. We found that while both reporter proteins were produced in cells at permissive temperatures, when cultures were shifted to 37°C, nearly all *ded1-ts* cells lost GFP expression while the majority of cells retained mCherry signal. This validates that the *ded1-ts* mutation does not universally block translation, but severely impacts translation controlled by the *FKS1* 5’UTR. In contrast, WT cells shifted to 37°C maintained expression of both reporters (Fig. 2B). These results indicate the *FKS1-GFP* reporter is highly sensitive to Ded1-inhibition.

**Figure 2.**
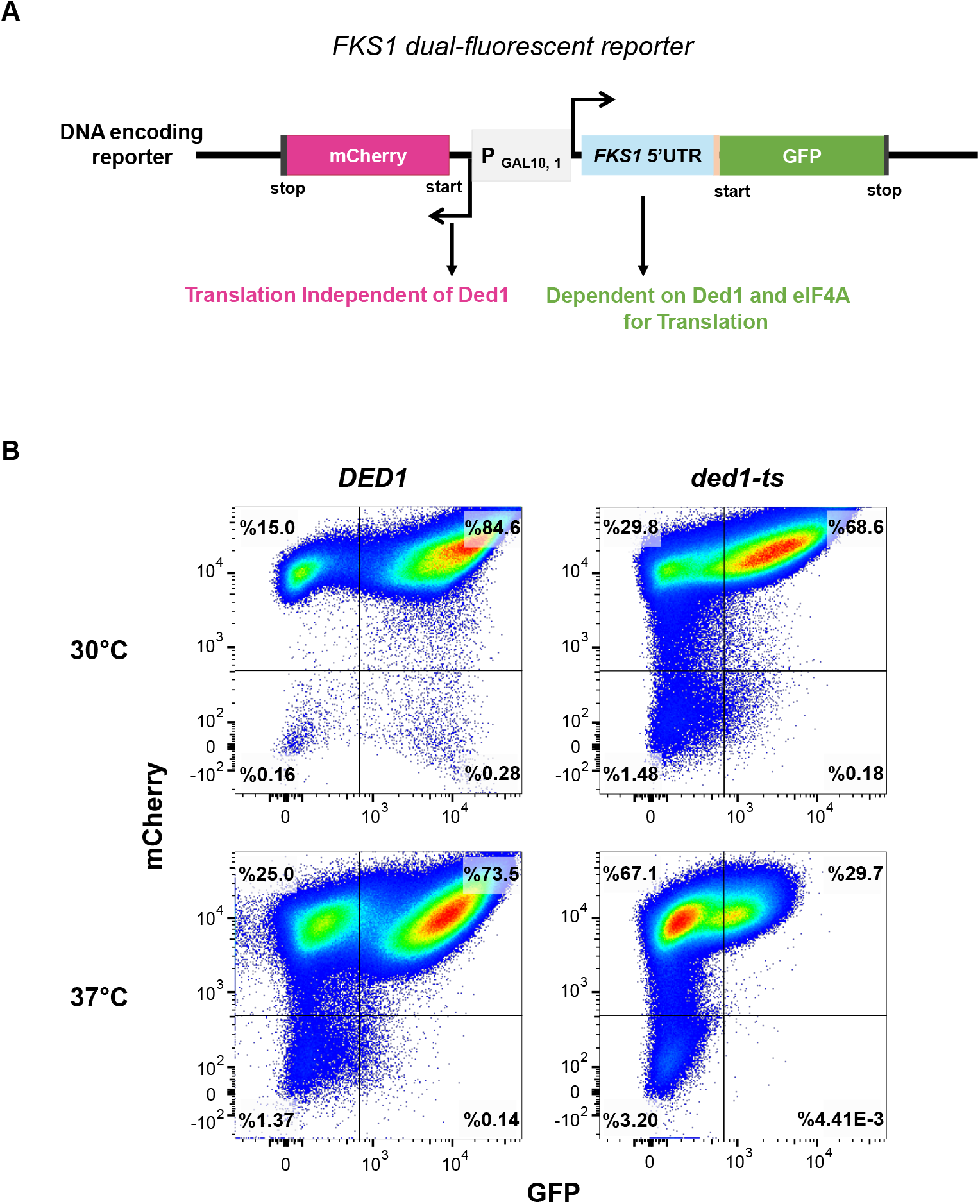
Validation of the *FKS1* fluorescent reporter for sensitivity to Ded1 inhibition. **A**. Schematic of the *FKS1* dual-fluorescence reporter construct, featuring a structured *FKS1* 52 UTR upstream of a superfolder GFP and an unstructured 52 UTR upstream of mCherry, driven by a bidirectional galactose promoter. **B**. Flow cytometry analysis of GFP and mCherry expression in *DED1* (YSW223) and *ded1-ts* (YSW222) strains grown at 30°C and 37°C to assess temperature-sensitive effects on translation of the reporter.

Next, the effects of combined RocA and caspofungin treatment on growth and translation were determined by growing reporter cells for 48 hours at 37°C with galactose induction, and monitoring OD_600_, GFP, and mCherry signals in an automated plate reader. WT and *ded1-ts* strains were treated with 0, 150, 200, or 500 μM RocA in combination with 0, 0.02, 0.04, or 0.06 μg/ml caspofungin (Fig. 3). While the suboptimal (0.02 μg/ml) caspofungin dose alone did not substantially inhibit WT growth, addition of 150 or 200 μM RocA with 0.02 μg/ml Caspofungin inhibited growth to a slightly higher degree than either drug alone (Fig. 3A left panel). This effect was further amplified with higher levels of caspofungin (Figure 3B and 3C, 0.04 and 0.06 μg/ml caspofungin). With the combination of 0.06 μg/ml caspofungin and either 150 or 200 μM RocA, WT cells exhibited almost complete growth inhibition (Fig. 3C). Additionally, 500 μM RocA alone nearly abolished growth (Fig. 3A-C, left panels), but combination of 0.06 μg/ml caspofungin and 500μM RocA abolished growth. Together, these results indicate that the RocA and caspofungin combination has higher efficacy than either drug on its own.

**Figure 3.**
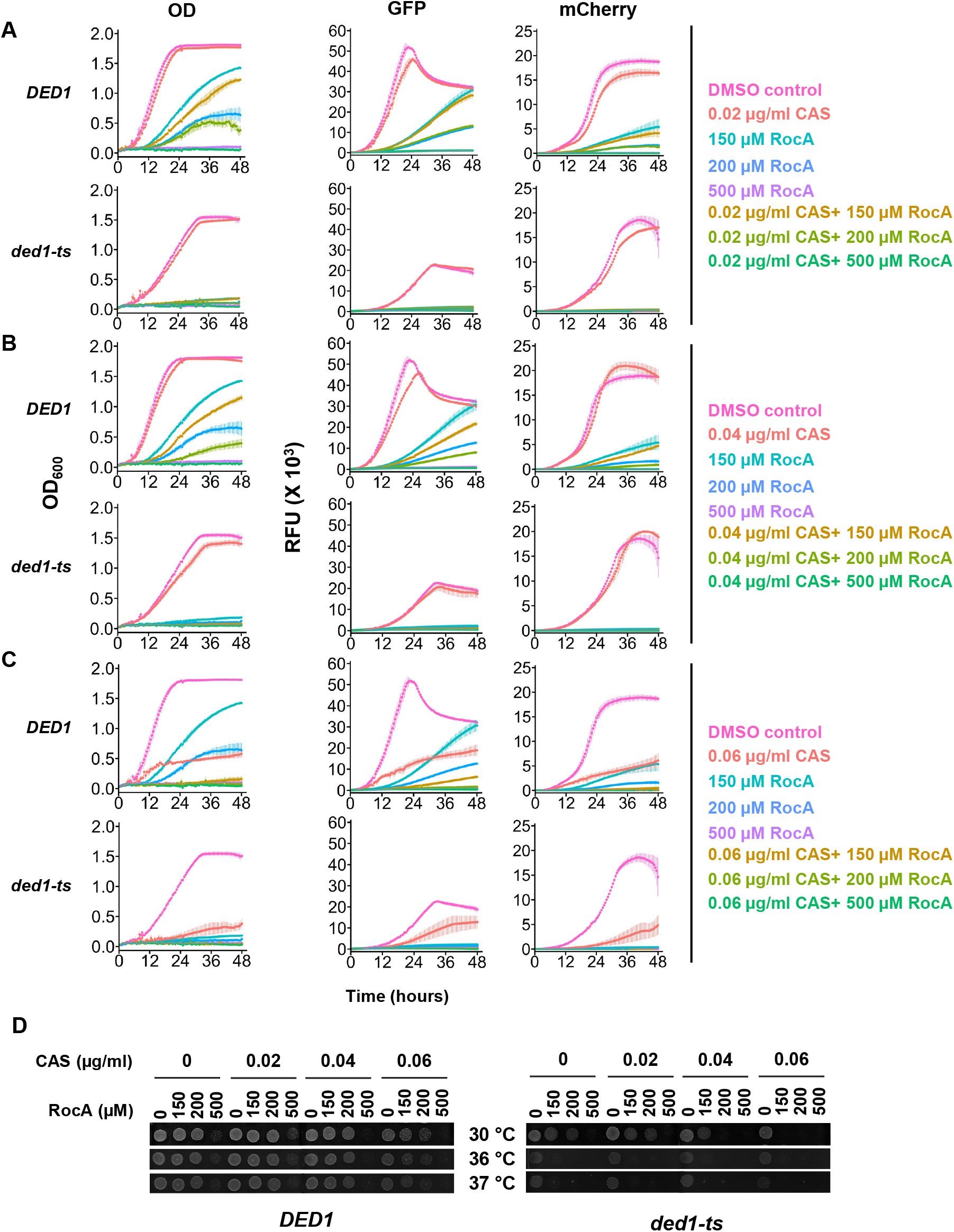
RocA combined with caspofungin led to enhanced inhibition of both growth and translation. **A-C**. 48-hour growth curves of *DED1* (YSW223) and *ded1-ts* (YSW222) strains treated with 150 μM, 200 μM or 500 μM RocA in combination with increasing concentrations of caspofungin: 0.02 μg/ml (A), 0.04 μg/ml (B), and 0.06 μg/ml (C). Controls included 1% DMSO (no drug), RocA alone, and caspofungin alone. The OD_600_, GFP, and mCherry signals were measured every 30 minutes at 37 to assess growth and reporter activity. Data represents the mean± standard deviation from 3 biological replicates. **D**. Recovery following combined RocA and caspofungin treatment: *DED1* and *ded1-ts* cells were spotted on YPD plates and incubated for one day at the indicated temperatures following indicated 48-hour treatments.

Next, we analyzed the effects of the drugs when Ded1 was inactivated by following growth of the temperature-sensitive mutant (*ded1-ts*). In this case, addition of 0.02 or 0.04 μg/ml caspofungin was tolerated on its own, but addition of any concentration of RocA, alone or in the presence of caspofungin, led to near-complete growth inhibition (Figure 3A and B, compare upper and lower OD panels for all but red and pink traces). This is expected given that RocA and the *ded1-ts* mutation target the same process to inhibit growth. We likewise found that with the highest concentration of caspofungin, 0.06 μg/ml, the *ded1-ts* mutant grew at much slower rates than WT cells, indicating caspofungin sensitivity is also impacted by Ded1 function.

We then analyzed effects on translation of the GFP reporter harboring the long and structured *FKS1* 5′ UTR and the mCherry reporter with a short unstructured 5′ UTR. The GFP and mCherry signals followed trends similar to OD_600_ measurements, consistent with a general defect in translation likely due to the inhibition of one or both helicases, Ded1 and eIF4A (Fig. 3A-3C). However, the *ded1-ts* strain showed ∼2-fold lower peak GFP levels in the absence of drug (Fig. 3A, 3B and 3C), illustrating that the Ded1 substitution has a strong inhibitory effect on *FKS1* translation even without addition of drug. A very modest effect of the Ded1 mutation was also observed for mCherry without drug or with the lowest caspofungin concentration, manifesting as a rate defect, and is consistent with Ded1 having both general effects on translation of all RNAs and specific stronger effects on translation of highly structured mRNAs (14, 15).

In the recovery assay, WT cells showed substantial growth after treatment with lower concentrations of caspofungin and RocA when recovered without drugs at all three temperatures (Fig. 3D). Cells exposed to higher concentrations of caspofungin in combination with lower doses of RocA also showed partial recovery under all temperatures. In contrast, treatment with the highest RocA concentration tested (500 μM) alone resulted in minimal recovery at each temperature, indicating strong but non-lethal growth inhibition. However, combining 500 μM RocA with higher concentrations of caspofungin was fungicidal, indicating the drug combination could prevent cells from developing new resistance mutations. For recovery of *ded1-ts* cells, synthetic lethality was observed. With 0.04 μg/ml caspofungin, or with 200 μM RocA alone, there was some recovery, but when the two drugs were combined at these concentrations, the cells did not recover, even at 30°C. Recovery of the mutant was modest in all cases, consistent with the slow growth of the mutant seen at 37°C, and suggesting most cells were inhibited by the elevated temperature incubation even without drug treatment. However, at 36°C and 37°C, *ded1-ts* cells could not recover from RocA treatment, suggesting targeting Ded1 specifically sensitizes cells to RocA and/or the combination therapy, particularly with temperatures relevant to human infection (Fig. 3D).

### RocA is an effective inhibitor of the fungal pathogen *Candida glabrata*

After observing synthetic effects of RocA with caspofungin on *S. cerevisiae* growth and translation, we extended our investigation to a pathogenic yeast to explore possible therapeutic applications. We selected *C. glabrata*, a species phylogenetically closer to *S. cerevisiae* than other *Candida* species (16). Moreover, because *C. glabrata* is resistant to certain antifungal drugs, new treatment strategies could be particularly impactful. We found that caspofungin inhibited growth of *C. glabrata* in a dose-dependent manner between 0.03 and 0.2 μg/ml caspofungin at its optimal growth temperature of 37°C, and cells were able to recover from these treatments (Fig. 4A and 4F). However, 0.3-0.5 μg/ml caspofungin completely blocked growth and prevented recovery (Fig. 4A and 4F). RocA also showed a dose-dependent inhibition of growth, but cells recovered well at all concentrations and temperatures tested (Fig. 4B and 4F).

**Figure 4.**
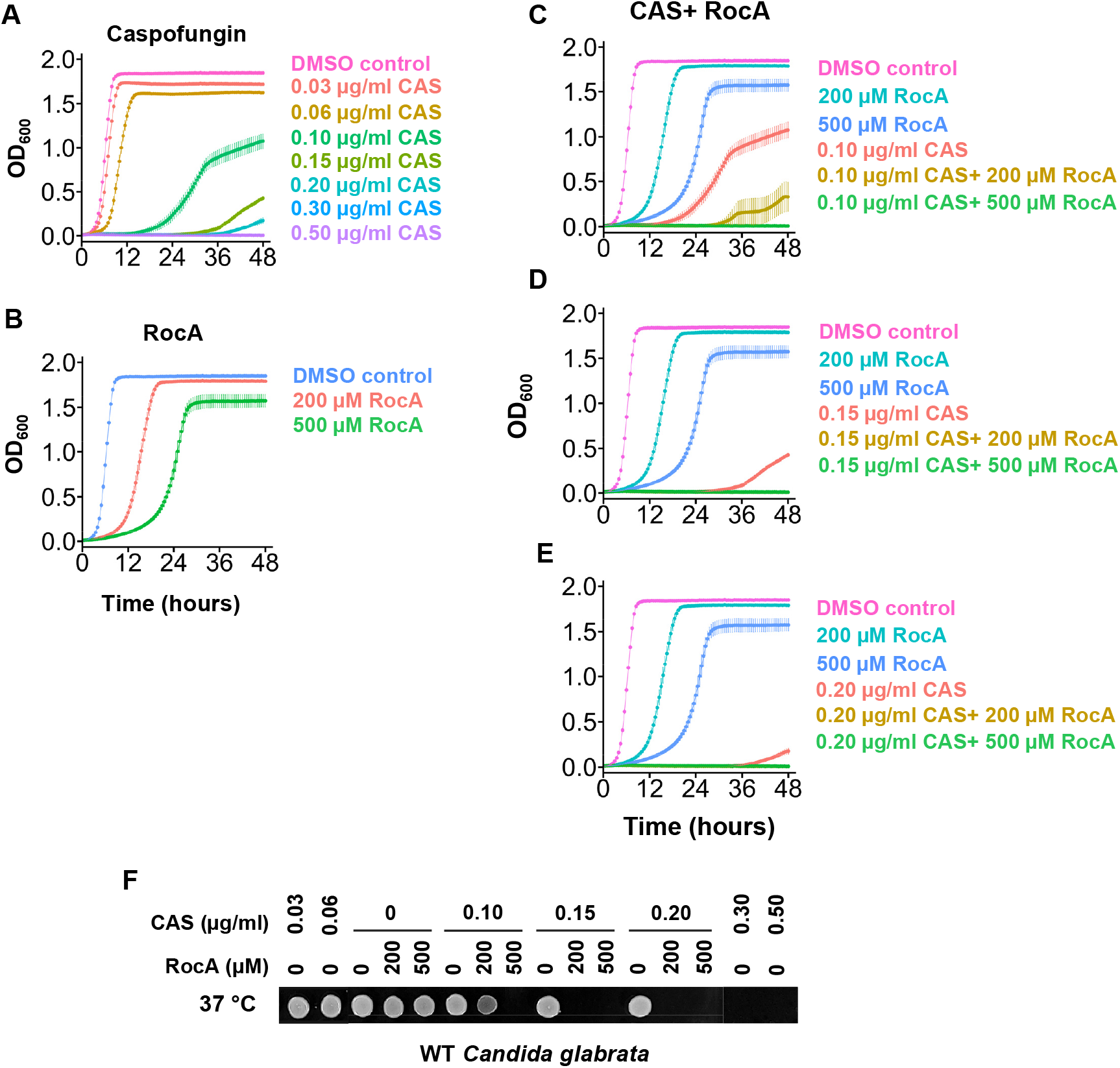
Effects of caspofungin, RocA and combinations in WT *C. glabrata*. **A-B**. 48-hour growth curves of WT *C. glabrata* treated with increasing concentrations of caspofungin (A), or RocA (B) at 37°C. OD_600_ was measured to monitor growth. Data represents the mean± standard deviation from 3 biological replicates. **C-E**. 48-hour growth curves of WT *C. glabrata* BG2 strain treated with 150 μM, 200 μM or 500 μM RocA in combination with increasing concentrations of caspofungin: 0.10 μg/ml (C), 0.15 μg/ml (D), and 0.20 μg/ml (E). Controls included 1% DMSO (no drug), RocA alone, and caspofungin alone. OD_600_ was measured every 30 minutes to assess growth at 37°C. Data represents the mean± standard deviation from 3 biological replicates. **F**. Recovery following compound exposure: WT *C. glabrata* cells were spotted on YPD plates and incubated at 37 °C for one day.

We then tested combinations of RocA and caspofungin on *C. glabrata* growth and recovery (Fig. 4C-F). WT *C. glabrata* cells were treated at 37°C with 0, 200, or 500 μM RocA in combination with 0, 0.10, 0.15, or 0.20 μg/ml caspofungin. Caspofungin alone at the tested concentrations inhibited growth more strongly than RocA alone, and its combination with either 200 μM or 500 μM RocA further reduced growth compared to either drug alone (Fig. 4C-E). In recovery assays, 200 μM RocA combined with 0.10 μg/ml caspofungin impaired recovery more than 0.10 μg/ml caspofungin alone (Fig. 4F). Furthermore, combining 200 μM RocA with higher concentrations of caspofungin, or 500 μM RocA with any tested concentration of caspofungin, completely abolished recovery (Fig. 4F). These results suggest that RocA, when combined with caspofungin, is fungicidal in *C. glabrata*. Altogether, our findings indicate a synthetic interaction between RocA and caspofungin that enhances antifungal efficacy beyond either compound alone. This supports the potential of targeting translation in combination with cell wall synthesis to impair antifungal resistance from developing in *C. glabrata*.

## DISCUSSION

Here we assessed the efficacy of simultaneously targeting protein expression and the catalytic activity of Fks1 on fungal cell growth. We used RocA to target translation initiation helicases with the goal of reducing Fks1 translation, and caspofungin, an established antifungal that directly inhibits the Fks1 protein. We observed that RocA alone or in combination with caspofungin significantly impaired both cell growth and translation of an *FKS1* reporter in both *S. cerevisiae* Ded1 WT and mutant strains, which could be due to inhibition of either or both primary translation initiation helicases, Ded1 and/or eIF4A (Fig. 1 and 3) (11). At higher concentrations of both drugs, cells did not recover after treatment, indicating the combination was fungicidal where individual drug treatments were fungistatic. These findings suggest caspofungin and RocA treatment could target the same process. In addition, *ded1-ts* cells showed even higher sensitivity to either drug or the combination, and addition of RocA was fungicidal at the infection-relevant restrictive temperature (Fig. 3D). This suggests global translation inhibition through RocA synergizes with specific inhibition of Ded1 to impair yeast growth and further sensitize cells to caspofungin. Both eIF4A and Ded1 were previously shown to enhance translation of *FKS1* and related genes, so it is possible that this is through *FKS1* expression, or through a more general translation inhibition mechanism. *FKS1* is functionally redundant with the paralog *GSC2* (17). Interestingly, this RNA was also observed to be dependent on Ded1 for translation, so it is possible that both β-1,3-glucan synthase RNAs show lower translation in cells treated with RocA or when Ded1 is impaired. Alternatively, it is possible that by repressing Fks1 expression, the amount of caspofungin required to obstruct Gsc2 is reduced.

We then extended our study to the yeast pathogen *C. glabrata* (Fig. 4). Previous research has shown that species sensitive to rocaglates, including *S. cerevisiae* and *C. auris*, contain a phenylalanine at the equivalent position of residue 151 in *S. cerevisiae* (10). *C. glabrata* has a phenylalanine at this position, yet while RocA alone inhibited growth at high concentrations, it was not fungicidal at the highest concentrations tested. This observation is consistent with previous reports showing that *C. glabrata* was less sensitive to the rocaglate compound CMLD010515 (RHT) than *S. cerevisiae* and several other *Candida* species (10). Interestingly, the addition of caspofungin overcame this limited sensitivity, possibly by compromising cell wall integrity and thereby enhancing RocA uptake and efficacy. The combination markedly reduced the concentration of caspofungin required to kill *C. glabrata* beyond an additive effect of the two drugs, further indicating a synergistic interaction.

In this work we used the commercially-available rocaglate, RocA, but other rocaglamides have recently been shown to have greater efficacy in hindering yeast growth (10). Our work suggests that combining these more inhibitory forms with caspofungin could allow use of lower concentrations of both drugs while maintaining desired fungicidal activity. RocA broadly hampers translation in mammalian cells, which could cause undesired side-effects, so reducing the concentration needed for antifungal therapy would benefit patients. We observed that a *ded1-ts* mutant specifically impaired Fks1 translation without affecting translation of an unstructured reporter, and showed synergy with RocA to inhibit growth. Given this observation, it is likely that specific inhibitors of Ded1 that do not also target eIF4A could further enhance sensitivity of fungal pathogens to caspofungin and RocA. RK-33, an inhibitor of the mammalian Ded1 ortholog DDX3 was shown to inhibit Ded1 in reconstituted unwinding assays, making it a promising prospect (18). However, it did not constrain yeast growth in our experiments, suggesting RK-33 is unable to penetrate the cell wall (Figure S1). It is possible that other fungal pathogens with distinct cell wall architecture could be more susceptible, or it may be possible to improve uptake of an RK-33 derivative for antifungal treatment. Altogether these findings highlight the potential of combining translation and cell wall inhibitors to impede the development of drug resistance in fungal pathogens, which could lead to novel antifungal treatment strategies.

## Acknowledgments

The authors wish to thank members of their laboratories as well as Paul Cullen, John Panepinto, Laura Rusche, and Michael Yu for input on this work. This work was supported by NIH grants R00GM119173 and R01GM139977 to S.E.Walker and institutional funds from Roswell Park to J. Barbi. The Roswell Park Flow and Immune Analysis Shared Resource is supported by NIH/NCI Cancer Center Support Grant P30CA013696.

---

**Figure S1.**
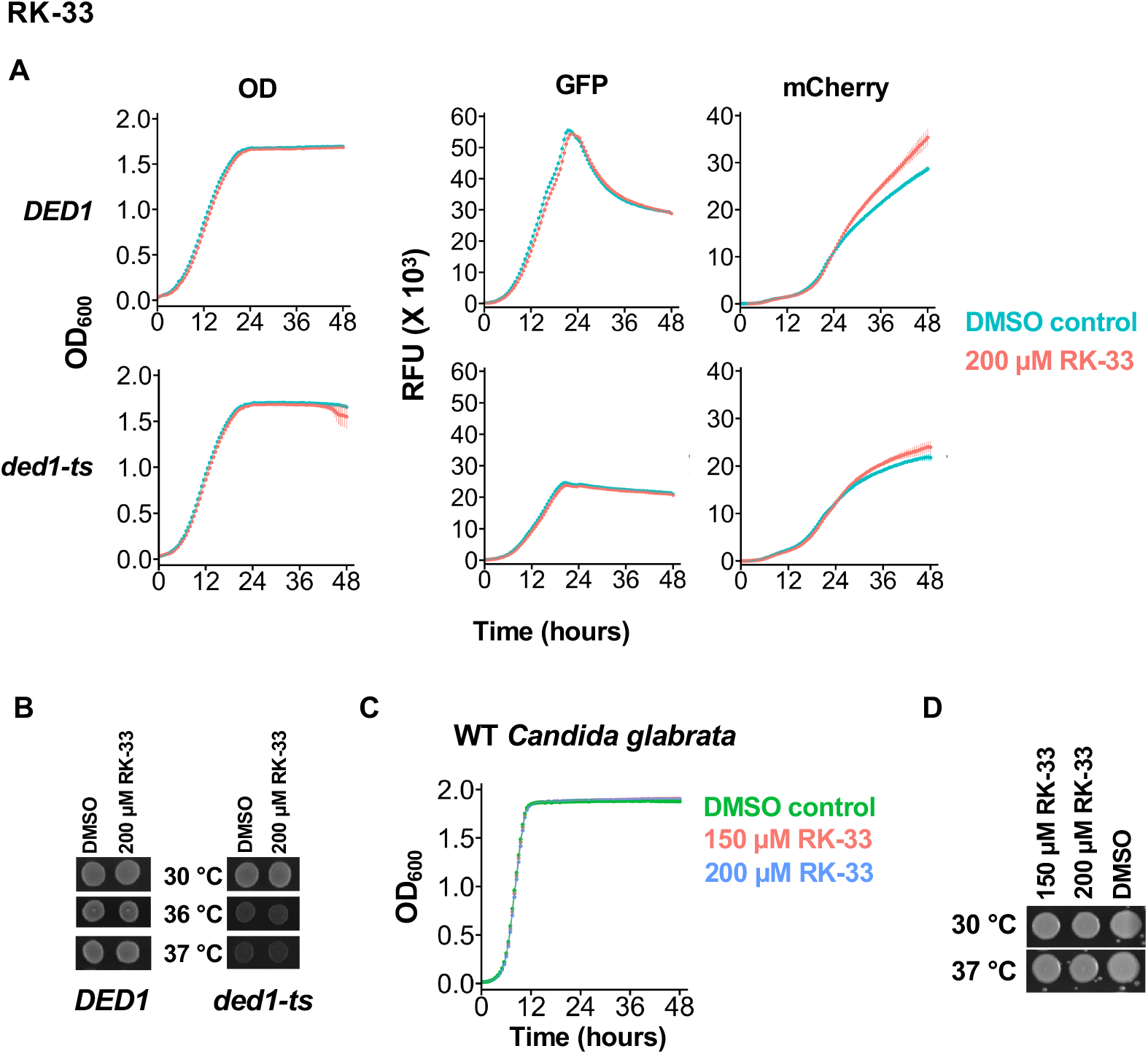
RK-33 was ineffective at inhibiting *S. cerevisiae* and *C. glabrata*. **A**. 48-hour growth curves of *DED1* (YSW223) and *ded1-ts* (YSW222) strains treated with 1% DMSO (no drug) or 200 μM RK-33 at 30°C. OD_600_, GFP, and mCherry signals were measured simultaneously to assess growth and reporter translation. Data represents the mean ± standard deviation from 3 biological replicates **B**. Recovery assay following DMSO and RK-33 treatment. *DED1* and *ded1-ts* cells (YSW223 and YSW222) were spotted on YPD plates and incubated for one day at the indicated temperatures to evaluate survival. **C**. 48-hour growth curves of WT *C. glabrata* treated with 0, 150 μM or 200 μM RK-33. OD_600_ was measured to monitor growth. Data represents the mean± standard deviation from 4 biological replicates. **D**. Recovery assay following compound exposure: WT *C. glabrata* cells were spotted on YPD plates and incubated at indicated temperatures for one day.

